# Divergent molecular phenotypes in point mutations at the same residue in beta-myosin heavy chain lead to distinct cardiomyopathies

**DOI:** 10.1101/2023.07.03.547580

**Authors:** Sarah J. Lehman, Artur Meller, Shahlo O. Solieva, Jeffrey M. Lotthammer, Lina Greenberg, Stephen J. Langer, Michael J. Greenberg, Jil C. Tardiff, Gregory R. Bowman, Leslie Leinwand

## Abstract

In genetic cardiomyopathies, a frequently described phenomenon is how similar mutations in one protein can lead to discrete clinical phenotypes. One example is illustrated by two mutations in beta myosin heavy chain (β-MHC) that are linked to hypertrophic cardiomyopathy (HCM) (Ile467Val, I467V) and left ventricular non-compaction (LVNC) (Ile467Thr, I467T). To investigate how these missense mutations lead to independent diseases, we studied the molecular effects of each mutation using recombinant human β-MHC Subfragment 1 (S1) *in vitro* assays. Both HCM-I467V and LVNC-I467T S1 mutations exhibited similar mechanochemical functions, including unchanged ATPase and enhanced actin velocity but had distinct effects on the basal activity of myosin. HCM-I467V S1 showed no change in basal ATPase activity of myosin while LVNC-I467T reduced the basal ATPase activity by 50%. Molecular dynamics simulations reveal that I467T allosterically disrupts nucleotide binding of myosin, which may contribute to the uncoupled reduced basal activity and enhanced actin velocity observed in this mutation. These contrasting molecular effects may lead to contractile dysregulation that initiates LVNC-associated signaling pathways that progress the phenotype. Together, analysis of these mutations provides evidence that phenotypic complexity originates at the molecular level and is critical to understanding disease progression and developing therapies.

## Introduction

The most common genetic cardiomyopathy is Hypertrophic Cardiomyopathy (HCM), typically characterized by thickened ventricular walls and impaired relaxation. This disease affects between 1 in 200 to 1 in 500 people in the United States and is the most common cause of sudden cardiac death in people under 30 years old. A leading cause of HCM are mutations in sarcomeric protein genes, and mutations in MYH7 (also known as cardiac myosin heavy chain; β-MHC) account for ∼30% of HCM[1, 2]. Until recently, HCM lacked any pharmaceutical intervention that targeted the cause of disease with existing treatments addressing symptoms of the disease by improving cardiac relaxation[3]. However, FDA approval of Camzyos©, a myosin-inhibitor, provided the first sarcomere-targeted therapeutic for the treatment of obstructive HCM[4]. Recently, a second-generation myosin inhibitor, Myqorzo, received FDA approval for treatment of obstructive HCM[5], highlighting the benefit of directly modulating sarcomere function to treat this complex disease.

Another form of cardiomyopathy that can be caused by mutations in MYH7 is left ventricular non-compaction (LVNC)[6-8]. In contrast to HCM, LVNC is a much less understood and studied disease. It is characterized by abnormal growth of cardiac tissue, with excessive deposition but reduced compaction of trabeculae, resulting in a non-compacted, spongiform-like myocardial wall (for review[9]). The introduction of cardiac MRI in the clinic improved the identification of this non-compacted phenotype from thickened, dense muscular wall commonly seen in HCM. While this morphology is characteristic of the disease, the functional phenotype is highly variable. LVNC may present with preserved cardiac function, systolic dysfunction, diastolic dysfunction, or a combination of both[10]. Thus, clinical management of disease is dependent on the clinical manifestation of the LVNC phenotype. Despite these variable presentations, the natural history of disease has been frequently associated with arrhythmias, heart failure, cardiac transplants, and risk of sudden cardiac death. Additionally, due to the spongiform-like myocardium, patients are at risk of thromboembolic events and typically require blood thinners as preventative treatment[11, 12].

Though mutations in the *MYH7* gene have been identified as the cause of disease in many cases, there still is no consensus on how mutations lead to specific cardiomyopathies. Numerous mutations, including those associated with HCM, have been shown to alter the mechanochemical function of myosin, typically enhancing ATPase activity of the motor domain[13-15]. Other mutations increase the number of myosin heads that are available to bind to actin and form force-generating cross bridges[16-18]. This dysregulation of the relaxed states of myosin may be driven by structural rearrangements of myosin heads[19-23] and/or biochemical function of myosin[13, 24-26]. Specifically, myosin is known to exist in two distinct states known as the actin-available, disordered (DRX) state and the energy-conserving, super-relaxed (SRX) state of myosin[24, 27, 28]. Multiple studies have shown a mutation-specific and disease-dependent shift in the SRX/DRX ratio, providing an additional biophysical mechanism for altered contractility and energetic expenditure in these cardiomyopathies[17, 21, 24]. In contrast to HCM, the molecular mechanisms that underlie the complex morphogenetic remodeling and altered cardiac function in LVNC are not well understood. Of note, one study investigated the primary insults caused by LVNC-associated β-MHC M531R, describing an increased power output driven by enhanced mechanochemical activity of the motor[29], despite reduced contractile performance reported in patients expressing this mutation[30].

Here, we investigate how two missense mutations at the same position in the β myosin heavy chain lead to two different diseases. Specifically, we investigate the biochemical mechanisms that govern how I467V (Ile467Val) causes HCM while I467T (Ile467Thr) causes LVNC. We performed a variety of biochemical assays that probed how these mutations affect intrinsic enzymatic function, actin sliding, and myosin-based relaxation properties. We also connect our experimental results to molecular dynamics simulations of the β-myosin motor domain carrying the I467T mutation to understand how a small structural perturbation can alter emergent ensemble properties that give rise to biochemical differences.

## Results

### Actin-Activated ATPase Activity

The enzymatic function of cardiac muscle contraction is driven by the mechanochemical activity of actomyosin interactions. To assess whether these similar point mutations alter the enzymatic function of the myosin motor, we performed an NADH-coupled actin-activated ATPase assay to measure maximal ATPase activity and actin affinity (Figure 1). The maximum ATPase activity of the HCM-I467V and LVNC-I467T S1 constructs were unchanged as compared to both β-MHC WT S1 (1.6 ± 0.32 s^-1^, 1.6 ± 0.57 s^-1^, and 1.5 ± 0.48s^-1^, respectively) (Table 1). The actin affinity for LVNC I467T S1 was increased, albeit not significantly, as compared to both β-MHC WT S1 and HCM-I467V S1 (48.2 ± 28.3 µM, 59.3 ± 31.1 µM, and 72.7 ± 33.6 µM, respectively). These data suggest that though these mutations are in close proximity to the ATP binding pocket, they do not alter the ATPase activity of the myosin motor domain.

**Table 1.** Summary of actin-activated ATPase activity of myosin S1 constructs.

|  | **β WT S1** |  | **β I467V S1** |  | **B I467T S1** |  |
| --- | --- | --- | --- | --- | --- | --- |
|  | Average | S.D. | Average | S.D. | Average | S.D. |
| **K<sub>cat</sub> (s<sup>-1</sup>)** | 1.5 | 0.48 | 1.6 | 0.32 | 1.6 | 0.57 |
| **K<sub>M</sub> (μM)** | 59.3 | 31.1 | 72.7 | 33.6 | 48.2 | 28.3 |

Data are represented as mean ± S.D. N=6-11 curves per myosin S1 construct.

**Figure 1.**
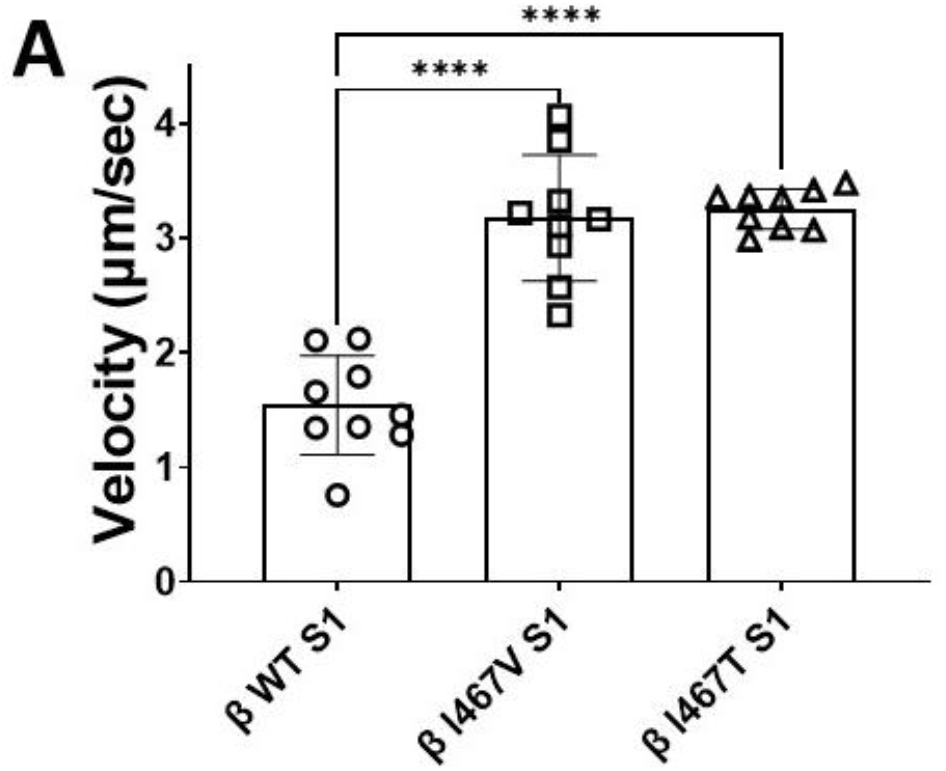
Actin-activated ATPase Activity of myosin constructs. ATPase activity of myosin S1 constructs was measured over time with increasing actin concentrations. Data points were fitted using Michaelis Menten kinetics to obtain maximum ATPase activity (k_cat_) and actin affinity (K_M_). N=6-11 curves per myosin S1 construct.

### Single Turnover ATPase Activity

A growing number of studies support the role of myosin in regulating the relaxation and activation of cardiac muscle. Using a modified single-turnover ATPase assay, we measured the activity of myosin in an actin-independent state to compare HCM-I467V and LVNC-I467T mutational effects on the relaxed state of myosin (Figure 2). HCM-I467V S1 slightly reduced the basal activity of myosin as compared to WT S1 (0.0094±0.0008s^-1^, 0.016±0.005 s^-1^ respectively, p= 0.011) (Figure 2B, Table 2). Interestingly, the I467T mutation reduced the basal activity more than 2-fold (0.0044±0.0004s^-1^, p<0.0001) (Figure 2B, Table 2). While both mutations increased the area under the curve (AUC, Figure 2C, Table 2), LVNC-I467T S1 had a near two-fold increase relative to HCM-I467V S1, suggesting a greater reduction of energetic expenditure in this mutation. Unlike the intrinsic ATPase activity described above, LVNC-I467T S1 markedly reduced the basal ATPase activity of myosin compared to HCM-I467V S1, suggesting that a primary driver for the distinct diseases may be regulation of myosin recruitment into an actin-activated state.

**Table 2:** Summary table of the single turnover assay of myosin S1 constructs.

|  | **β WT S1** |  | **β I467V S1** |  | **B I467T S1** |  |
| --- | --- | --- | --- | --- | --- | --- |
|  | Average | S.D. | Average | S.D. | Average | S.D. |
| **k (s<sup>-1</sup>)** | 0.016 | 0.005 | **0.0094\*** | **0.0008** | **0.0044\*\*\*\*** | **0.0004** |
| **Lifetime (sec)** | 68.8 | 22.2 | **106.5\*\*** | **8.9** | **227.2\*\*\*\*#** | **21.2** |
| **AUC** | 9615 | 2696 | 11887 | 1056 | **21528\*\*\*\*#** | 1438 |

**Figure 2.**
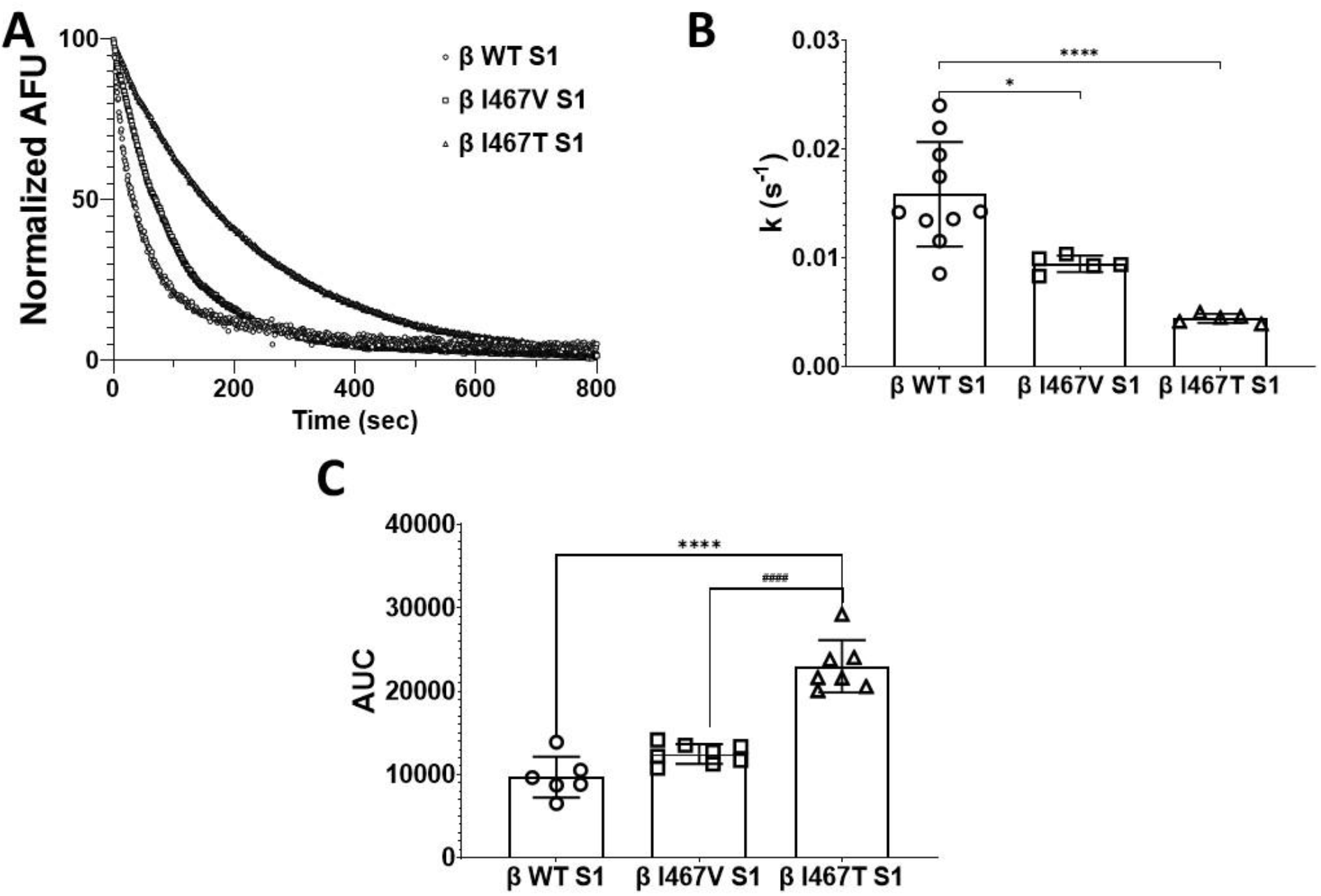
Single turnover ATPase assay of myosin S1. (A) Representative single-exponential fluorescence decay curves from the single-turnover mant-ATP dependent ATPase assay for myosin S1 constructs (WT: blue, β-I467V: red, β-I467T: green). (B) The basal ATPase activity was calculated from the single-exponential fit of decay curves. (C) Area under the curve (AUC) was calculated for each decay curve. A one-way ANOVA was used to assess statistical significance (* p = 0.0108, **p=0.0073, ****= p<0.0001 vs. β-WT S1) and adjusted with Tukey’s multiple comparisons to assess differences between mutations (# = p<0.0001 vs. β-I467V S1). Values are calculated from 5-10 decay curves per S1 construct.

### In vitro Motility

The velocity by which myosin moves actin is a highly characterized parameter to understand the effects of mutations on the mechanical function of myosin. To assess the effects of the HCM-I467V and LVNC-I467T mutations, we utilized an unloaded in vitro motility assay and measured the velocity of actin movement for each S1 mutation. HCM-I467V and LVNC-I467T S1 increased actin velocity >2-fold each, (HCM-I467V S1: 3.18 ± 0.55 µm/sec, p=0.0025 and β-I467T S1: 3.26 ± 0.18 µm/sec, p<0.0001) as compared to WT-S1 (β-MHC WT S1: 1.38 ± 0.33 µm/sec) (Figure 3, Table 3). These data suggest that despite these two mutations leading to distinct cardiomyopathies, they alter the mechanical function of myosin in a similar manner. In combination with the ATPase activity, these data suggest that HCM-I467V and LVNC-I467T do not uniquely alter the intrinsic function of the myosin motor or its ability to translocate actin; rather, these mutations cause distinct cardiomyopathies in a mechanochemical-independent manner, but dependent on regulation of myosin activation.

**Table 3:** Summary table for In vitro motility actin sliding velocity for myosin S1 constructs.

|  | **β WT S1** |  | **β I467V S1** |  | **B I467T S1** |  |
| --- | --- | --- | --- | --- | --- | --- |
|  | **Average** | **S.D.** | **Average** | **S.D.** | **Average** | **S.D.** |
| **Velocity**<br>**(μm/sec)** | **1.38** | **0.33** | **3.18\*\*\*\*** | **0.55** | **3.26\*\*\*\*** | **0.18** |
| **% Motile** | **79.3** | **7.8** | **79.5** | **9.8** | **77.7** | **11.8** |

Values are reported as mean ± S.D. A one-way ANOVA was used to assess statistical significance (\*\*\*\*= p<0.0001) compared to β-MHC WT S1. N = 7-8 videos per myosin S1 construct.

**Figure 3.**
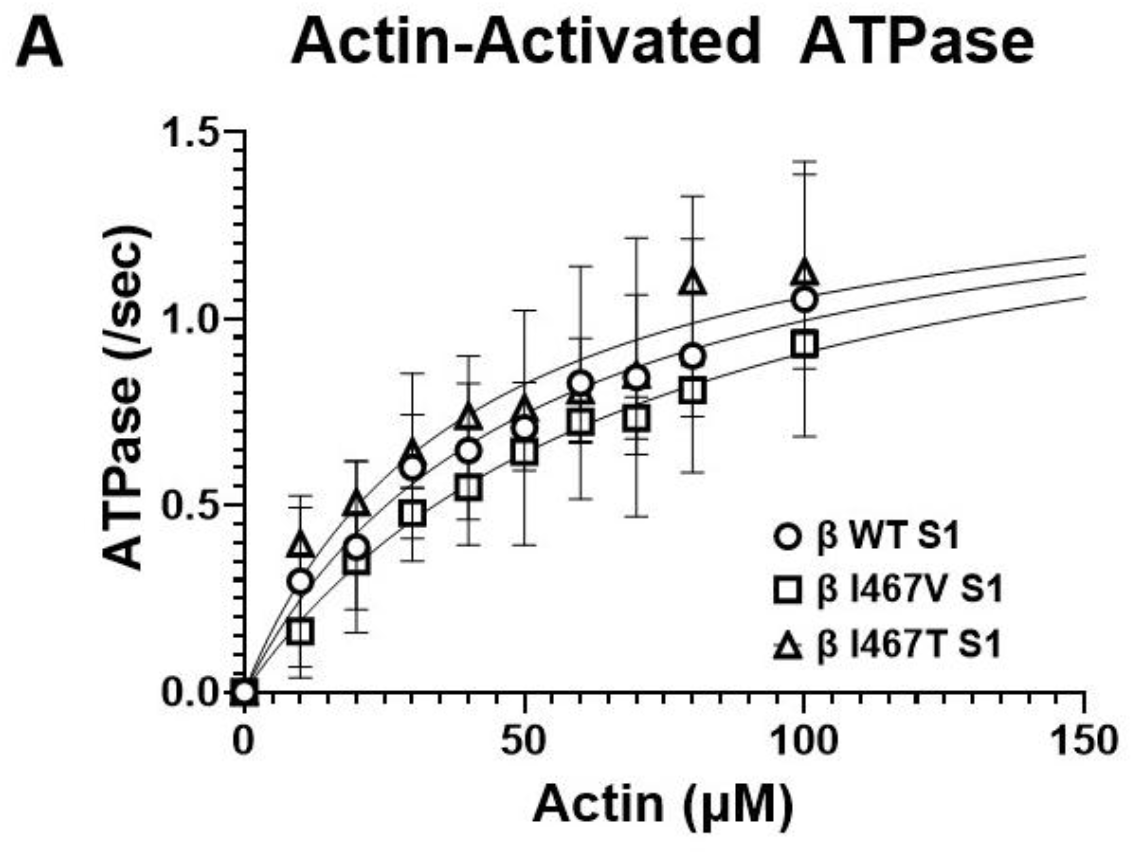
Unloaded Actin Sliding Velocity. In vitro motility rates for myosin S1 constructs were measured using TRITC-labeled actin filaments. Average actin sliding velocity was calculated from actin filaments that were ≥1 µm in length and mobile in >15 frames per video. A one-way ANOVA was used to assess statistical significance (****= p<0.0001) compared to β-MHC WT S1. N = 7-8 videos per myosin S1 construct.

### Molecular Dynamics Simulations of I467T

Differences in myosin’s sliding velocity can result from changes in its mechanical and/or enzymatic function. Compared to the HCM-I457V mutation, LVNC-I467T resulted in a dramatic uncoupling of the chemomechanical activity of β-cardiac myosin, as reported above. We sought to understand if this uncoupling of function was due to changes in myosin structure, specifically within myosin’s active site. The I467T mutation introduces a polar residue at a cluster of hydrophobic residues near the active site that we hypothesized would allosterically weaken interactions between the active site and ADP, thereby accelerating ADP release. In support of this hypothesis, previous studies have shown that mutations in Switch II can allosterically affect the nucleotide binding pocket in myosin V[31]. Therefore, we sought to test if a similar trend is observed in β-cardiac myosin that may facilitate faster ADP release.

We used molecular dynamics simulations to investigate the effects of I467T on β-cardiac myosin dynamics. We ran long simulations of wild-type human β-cardiac myosin and I467T starting from a homology model of a pre-powerstroke crystal structure (PDB ID: 5N6A) bound to ADP and Pi. We ran 8 trajectories with an average length of 1300 ns (aggregate time = 10.5 μs) for WT and 47 trajectories with an average length of 1000 ns (aggregate time = 47.7 μs) for I467T. To understand the effect of the mutation on ADP and Pi contacts, we calculated contact probabilities for ADP to the nucleotide binding pocket and for Pi to neighboring residues. Additionally, we calculated the distribution of distances between atoms near the mutation and within the nucleotide binding pocket.

Although I467T is in Switch II, we found that I467T allosterically disrupts interactions between ADP and the nucleotide-binding pocket. In nucleotide-bound crystal structures of β-cardiac myosin, the nucleotide’s base forms a hydrogen bond with Y134 (Figure 4B). In simulations of WT β-cardiac myosin, this hydrogen bond typically remains formed (p_contact_= 63.9%, Figure 4C). In contrast, in simulations of I467T, this hydrogen bond is more likely to be broken than formed (p_contact_= 6.6%, p < 0.001, Figure 4C). Furthermore, in nucleotide-bound crystal structures of β-cardiac myosin, there are two residues, W130 and V186, that form a lid above the nucleotide’s base (Figure 4D), likely preventing the nucleotide from leaving the active site. Our simulations suggest that the interaction between W130 and V186 is more likely to break in the I467T than in the WT ensemble. The contact probability for W130 and V186 is 19.4% in WT but only 0.7% for I467T (Figure 4B, p < 0.001). Simulations of I467T sample partial dissociation events where the ADP is far from its crystal pose (e.g., ADP-N6 to Y134-OH distances of over 1 nm, Figure 4C). Taken together, these results suggest that ADP is not as tightly bound in the nucleotide binding pocket of β-cardiac myosin harboring the I467T mutation, providing a structural mechanism for the changes in *in vitro* motility.

**Figure 4.**
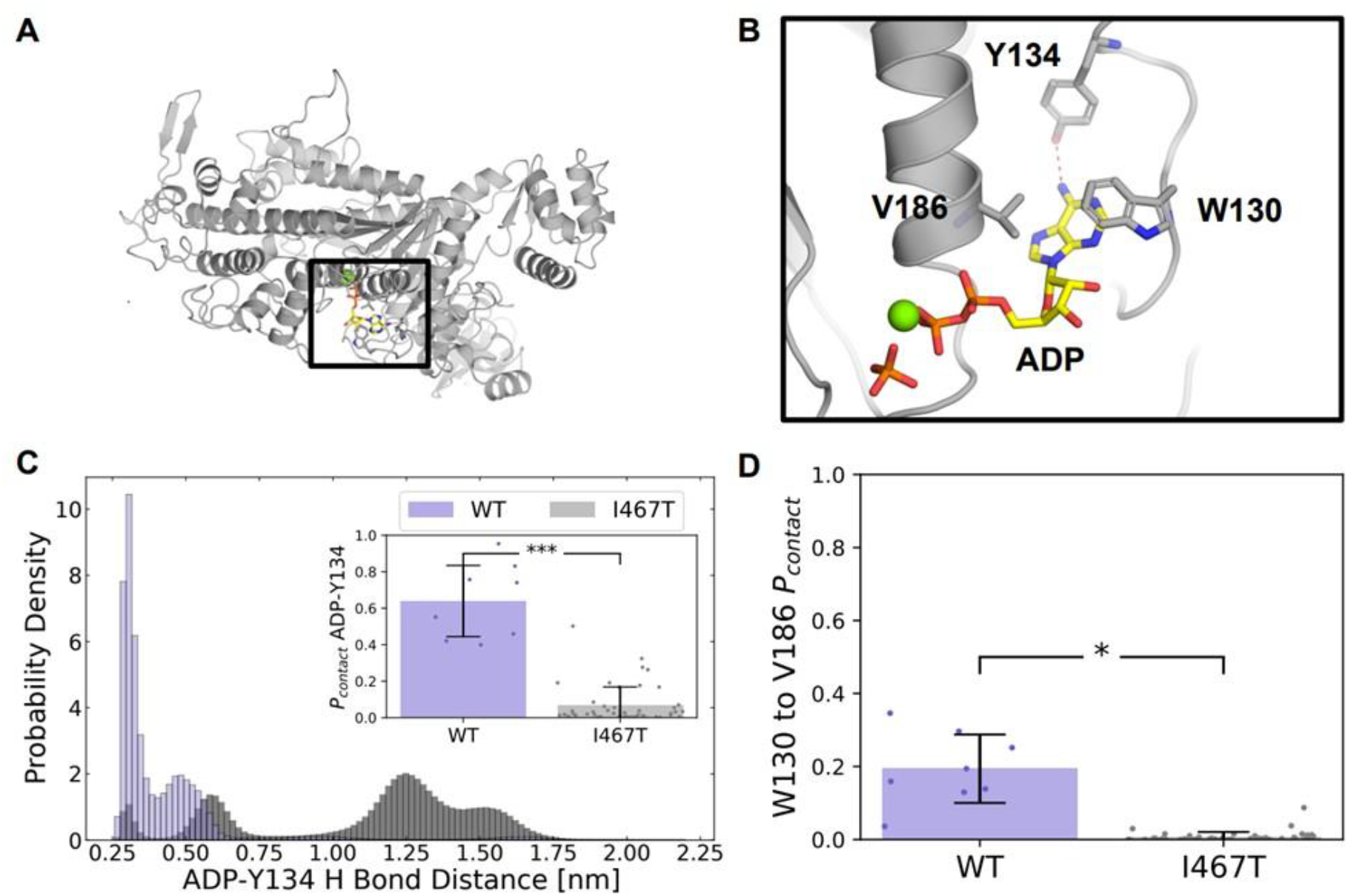
I467T disrupts interactions between ADP and the nucleotide-binding pocket in molecular dynamics simulations. (A) Homology model structure of human beta-cardiac myosin, (B) focused in on the ADP-Pi binding site, with ADP depicted in yellow and orange, Pi depicted in orange, and Mg2+ in green. The red dotted line indicates a hydrogen bond between adenine and Y134 that is seen in nucleotide-bound crystal structures. (C) I467T disrupts a native hydrogen bond between adenine (ADP-N6) and Y134’s hydroxyl group (Y134-OH). Additionally, the inset in (C) shows that the probability of a contact between ADP and TYR134 is significantly lower in I467T (p < 0.0001). (D) The contact probability between W130 and V186, two residues that form a lid over the adenine nucleobase, is significantly lower in I467T (p = 0.0011).

## Discussion

In this work, we investigated two similar missense mutations in β-myosin heavy chain: HCM-linked Ile467Val (I467V) and LVNC-linked Ile467Thr (I467T). To investigate how these mutations lead to unique pathologies, we studied the molecular effects of each mutation using recombinantly expressed myosin in various *in vitro* assays. Neither I467V nor I467T altered the maximal actin-activated ATPase activity (Figure 1), while both mutations showed a >2-fold increase in actin sliding velocity (Figure 3), respectively. Finally, I467T showed a significant reduction in basal myosin ATPase activity compared to WT and I467V (Figure 2). Thus, both I467T and I467V uncouple the canonical mechanochemical function of myosin described by the intrinsic enzymatic (ATPase) activity and the mechanical (actin sliding) properties of the motor, while only LVNC-I467T alters the regulation of the relaxation states of myosin.

While hundreds of sarcomeric mutations have been identified and linked to HCM[32, 33], the molecular mechanism by which these mutations converge on disease has not been identified. Historically, HCM was described as a gain-of-function disease as many mutations in β-MHC lead to enhanced mechanochemical function of myosin. HCM-I467V showed no change in the actin-activated ATPase activity, suggesting that the mutation does not alter the enzymatic function of the motor domain. However, HCM-I467V S1 did significantly increase the velocity by which myosin moves actin filaments. It is important to note that these assays are dependent on distinct steps of the mechanochemical cycle. The ATPase assay is an attachment limited assay in which phosphate release is the rate-limiting step, while the actin sliding assay is detachment limited, rate-limited by the release of ADP[34]. Though this mutation is located in the phosphate leaving tunnel of the myosin motor[35, 36], the unchanged k_cat_ reported for this mutation suggests that phosphate release is unchanged in this mutation, whereas the enhanced actin velocity in the HCM-I467V S1 mutation may be due to an increase in ADP release rate. Interestingly, molecular dynamic simulations by Li et al defined a link between the size of the phosphate tunnel and regulation of the SRX state[37], providing further evidence for primary structure-function disruption for the mutations studied in this work. However, direct investigation of the kinetics of ATP product release and SRX population effects were not investigated here and will be necessary to confirm mutation-specific changes in these parameters.

While the clinical manifestation of HCM is hypercontractile function, one of the primary drivers of disease is an inability to relax properly[38]. Recent studies have shown that many HCM-linked mutations in β-MHC disrupt the off-state of myosin, leading to an imbalance in the number of myosin heads available to bind actin during relaxation[17, 24, 25]. While the basal ATPase activity of myosin was slightly reduced with these mutations, the hypercontractile phenotype associated with HCM may be due to a shift toward the DRX state of myosin, supporting the increased velocity described above. However, the S1 protein used in this study is not sufficient to assess these population differences[39, 40] and thus was not assessed here.

LVNC-I467T S1 exhibited similar mechanochemical function to the HCM-I467V S1 mutation, including unchanged total actin-activated ATPase cycle time (the inverse of k_cat_) and enhanced actin velocity (Figures 1A and 3A). Computational modeling by Chakraborti et al suggested that I467T does not alter the rate of ATP hydrolysis into ADP and phosphate[35]. This is supported by the unchanged ATPase activity of I467T measured above. The magnitude of chemomechanical uncoupling seen in this mutation warranted further investigation into the structural changes that are caused by the mutation. Previous studies have shown that sequence variation in myosin motors modulates the probabilities of conformations primed for specific functional roles[41, 42]. Given the increased sliding velocity measured in the I467T mutation, we hypothesized that it would alter the structure of the ADP pocket myosin structure in a way that maintained the pocket would be in a more open state to facilitate ADP release. Though we did not observe complete ADP dissociation in simulations, we found that motors carrying the I467T mutation were more likely to adopt conformations primed for ADP release (i.e., structures with an open ‘lid’ and broken hydrogen bonds between ADP and myosin). This result strongly suggests that although the threonine does not directly interact with ADP, it allosterically disrupts interactions between ADP and the nucleotide-binding pocket.

Interestingly, Chakraborti et al. also found that I467T stabilized the post-rigor state of myosin, preventing the structural rearrangements necessary for the recovery stroke and pre-power stroke structure. While it is thought stabilization of the pre-power stroke state of myosin is required for increasing the ultra-slow SRX state of myosin, multiple studies have suggested additional SRX structural states that have not yet been identified[43-45]. Thus, one can imagine that by slowing the recovery stroke of myosin, I467T induces an undefined conformation that increases the ultra-slow state of myosin. These simulations are supported by the reduction in basal ATPase activity that was measured in LVNC-I467T S1 (Figure 2) and may indicate an increased SRX population in sarcomeres with this mutation. Additionally, the family affected by this mutation presents clinically with reduced systolic function, suggesting that the LVNC-linked I467T initiates disease, in part, by reducing myosin function and possibly reducing the availability of myosin heads for force-generation. This complex combination of altered mechanochemical function coupled to the reduced number of heads available to interact with actin may trigger a downstream signaling pathway distinct from both HCM and dilated cardiomyopathy, leading to the development of LVNC.

Development of non-compaction has been linked to unregulated Notch signaling which is sufficient to induce hypertrabeculation. Interestingly, one study showed that single molecule forces of around 4 pico-Newtons are sufficient to induce Notch signaling[46], forces that are within the range produced by a single molecule of cardiac myosin[47]. As *in vitro* motility reports on myosin in the actin-bound state, it is possible that the faster cycling of actin-bound I467T myosin could increase cardiac strain, resulting in activation of Notch signaling and enhanced trabeculation. However, because fewer heads are available to bind actin as described above, the inconsistent mechanoactivation may disrupt the normal signaling pathway, resulting in improper compaction. Additional investigation is necessary to understand the cellular signaling disruptions that arise in the presence of I467T that lead to the non-compaction phenotype.

While further studies are required to define the precise mechanisms by which these mutations lead to complex cardiac diseases, our work suggests that phenotypic complexity originates at the molecular level. Thus, a molecular understanding is critical to developing targeted therapies to treat discrete diseases that arise from similar point mutations. By directly modulating the molecular progression of disease, myosin- and/or sarcomere-targeted small molecule therapies offer a path to treat many patients.

## Methods

### Recombinant Myosin Construct Generation and Protein Purification

Human recombinant β myosin S1 constructs (residues 1-842) were used to assess the functional consequences of mutations located within the motor domain. Site-directed mutagenesis was performed using the Agilent Quikchange II XL kit (Agilent Technologies, Santa Clara, CA) was used to individually introduce the I467V and I467T mutations into the S1 motor domain and confirmed via cDNA sequencing. Constructs were tagged with a C-terminal affinity tag complementary to the PDZ binding domain [48]. Viral construct, protein generation, and protein purification were performed as described by Lee et al [49]. Tissue purified rabbit skeletal actin was also purified as previously described [49].

### ATPase Assay

Steady-state actin activated ATPase rates were measured using purified β-MHC S1 fragments expressing either the I467V or I467T mutations using an NADH-coupled regenerative ATPase assay. Purified wild-type (WT) β-MHC S1 was used as a control in all experiments. β-MHC S1 (f.c. = 0.4µM) was mixed with filamentous actin (concentration range: 0-100 µM) in the following solution: 20mM HEPES pH 7.0, 25mM KCl, 5mM MgCl_2_, 2mM ATP, 5mM DTT, 3mM phosphoenol pyruvate, 1mM NADH, 0.8mM pyruvate kinase/lactate dehydrogenase. Absorbance of NADH was monitored at 340nm over 60 minutes as a surrogate for ATP hydrolysis by the recombinant myosin. Linear absorbance ranges were fitted with Michaelis-Menten kinetics using GraphPad Prism 9 software. Basal ATPase activity (0 µM actin) was subtracted from ATPase rates were reported as k_cat_ (s^-1^) and actin affinity reported as K_m_ (µM).

### Single Turnover ATPase Assay

In order to assess the basal myosin ATPase activity, purified recombinant S1 proteins were run in a modified ATPase assay. To deplete ATP from the myosin S1 proteins, proteins were individually dialyzed exhaustively against a buffer containing: 10mM Tris pH 7.4, 30mM Potassium Acetate, 1mM EDTA, and 4mM MgCl_2_. Myosin S1 (f.c. = 0.4µM) was rapidly mixed with 0.4µM mant-ATP (2’-(or-3’)-*O*-(*N*-Methylanthraniloyl) Adenosine 5’-Triphosphate) and the reaction aged for 60 seconds to allow for mant-ATP + S1 binding and ATP hydrolysis. The reaction was then rapidly mixed with 4mM Na-ATP and decay of mant-ATP signal was immediately monitored using a BMG ClarioSTAR plate reader (BMG Labtech, North Carlina, USA). Specifically, mant-ATP was excited at 385nm and emission was monitored at 450nm at 1 Hz for 1000 seconds. The normalized decay curves were fitted with a single-exponential decay (constrained fit of Y_0_=100 and plateau = 0) using GraphPad Prism 9.0. The ATPase rate (K, s^-1^) and lifetime were calculated for β-MHC I467T and I467V and compared to β-MHC WT. The area under the curve was also calculated for all constructs to assess ATP consumption.

### Molecular Dynamics Simulations

A homology model of beta-cardiac myosin in the pre-powerstroke state from Meller et al. was used as the starting structure for simulations of WT and I467T.

All simulations were performed in the GROMACS simulation software with the CHARMM36m force field [50, 51]. Each simulation was prepared by solvating the myosin motor domain with TIP3P water in a dodecahedron box extending 1 nm around the protein. The system was then neutralized and raised to a salt concentration of 0.1 M NaCl. Energy minimization was performed using steepest descent till the maximum force on any atom was below 1000 kJ/(mol x nm). Equilibration was performed using the Bussi-Parrinello thermostat and the Parrinello-Rahman barostat with all heavy atoms restrained at 300 K.

Production runs were performed at 310 K with the leapfrog integrator, Bussi-Parrinello thermostat, and Parrinello-Rahman barostat. A 12 Å cutoff distance was utilized with a force-based switching function starting at 10 Å. We employed periodic boundary conditions to mimic the effects of bulk solution. The PME method with a grid density greater than 1.2 Å^3^ was used to calculate long-range electrostatics. All hydrogen bonds were constrained with the LINCS algorithm to enable the use of a constant integration timestep of 2 fs. All simulations were performed on our in-house compute cluster consisting of NVIDIA A5000 nodes.

### Simulation Analysis

We used MDTraj’s contacts function, which takes a subset of residues and reports any contacts between their atoms if any two atoms are within a cutoff threshold distance of each other; we defined the cutoff distance as 4 Å and used heavy atoms only. For each trajectory, we calculated the amount of time a contact exists between two residues and divided that by the length of the trajectory, yielding a probability of contact between those residues. The scripts for calculating contacts, statistical testing, and graphing the data can be found here: https://github.com/ssolieva/i467t.

### In vitro Motility

*In vitro* motility was used to assess the effect of I467T and I467V mutations on myosin’s ability to translocate actin. Purified recombinant S1 mutants and WT proteins were mixed with filamentous actin in a 1:3 ratio and 1mM ATP then subsequently spun at 90,000rpm for 25 minutes at 4°C to remove inactive myosin molecules. The active myosin S1 molecules were diluted to a final concentration of 0.35µM in a 1X buffer containing: 20mM HEPES pH 7.5, 25mM potassium chloride, 5mM MgCl_2_, 1mM EGTA, and 10mM DTT. Glass cover slips were dipped in 0.2% nitrocellulose and adhered to a glass microscope slide to create a flow-cell for the experiment. To begin, 3 µM SNAP-PDZ was flowed into the cell and incubated for 2 minutes at room temperature. 1mg/mL BSA was then added and incubated for one minute at room temperature to block all non-specific binding sites. 0.35µM myosin S1 solution was added to the flow chamber and incubated for three minutes to ensure proper binding to the surface mobilized SNAP-PDZ domain. Excess myosin was washed out, and phalloidin-labeled filamentous actin (f.c. 2-5nM) was added and incubated for one minute. Finally, activating solution containing 20mM HEPES pH 7.5, 25mM potassium chloride, 5mM MgCl_2_, 1mM EGTA, and 10mM DTT, 15mM ATP, 4mg/mL glucose, 0.135mg/mL glucose oxidase, 0.0215mg/mL catalase, and 0.5% methylcellulose was added to the flow chamber and incubated for 30 seconds at room temperature. Using an inverted wide-field microscope (Nikon Inc. New York, USA), movement of the phalloidin-labeled actin was monitored over time. Videos were collected between one and three frames per second and a total of 30 frames were collected for each movie. Using MATLAB, actin filament velocity was calculated for individual filaments and averaged across videos. At least 50 filaments were measured for each mutation, across three separately purified samples for β-MHC WT, I467T, and I467V S1.

### Statistical Analysis

For actin-activated ATPase assays, single turnover ATPase assays, *in vitro* motility assays, a one-way ANOVA was used to assess significant differences. Tukey’s multiple comparisons test was applied post-hoc to determine the mutation-specific effects in these assays. For simulation analysis, we conducted a student’s t-Test when the variances were approximately equal and Welch’s t-Test when the variances were unequal. For all comparisons, significance was assigned at p>0.05.

## Acknowledgements

We thank Ariana Combs for producing recombinant myosin protein. We thank Dr. Suman Nag for valuable discussions and insight. We thank Dr. Jian Wei Tay for motility analysis script generation and the BioFrontiers Institute Advanced Light Microscopy Core for use of microscopy equipment and for imaging support.

## Author Contributions

S. J. Lehman., J. C. T., and L. A. L. conceptualization; A. M., J. M. L., S. O. S., and M. J. G. methodology; J. W. T. software; S. J. Lehman., A. M., J. M. L., S. O. S., and L. G. formal analysis; S. J. Lehman., A. M., J. M. L., S. O. S., and L. G. investigation; A. M. and J. M. L., S. O. S. data curation; S. J. Lehman. and A. M. writing–original draft; S. J. L., A. M., J. M. L., S. O. S., S. J. Langer, M. J. G., J. C. T., G. R. B, and L. A. L. writing–review and editing; S. J. Lehman., A. M., J. M. L., S. O. S., and L. G. visualization; M. J. G., J. C. T., G. R. B., and L. A. L. supervision; M. J. G., J. C. T., G. R. B., and L. A. L. funding acquisition.

## Funding and additional information

This work was supported by the National Institutes of Health (NIH grant R01GM029090; to L. A. Leinwand), NIH training grant (grant no.: T32 HL007822-20; to S. J. Lehman), NIH grant R01HL141086; to M. J. G., and NIH grants R01GM124007 and RF1AG067194; to G. R. B.). A. M. was supported by the National Institutes of Health F30 Fellowship (1F30HL162431-01A1). This work was also supported by the National Science Foundation (NSF grant DGE-2139839 to J. M. L.; and NSF CAREER Award MCB-1552471; to G. R. B.) G. R. B. holds a Packard Fellowship for Science and Engineering from The David & Lucile Packard Foundation. The content is solely the responsibility of the authors and does not necessarily represent the official views of the NIH, the National Science Foundation, or the European Commission.

